# Dual role of the RIG-I 2CARD as a signaling module and a brake on dsRNA translocation

**DOI:** 10.64898/2026.10.13.758695

**Authors:** Sota Kondo, Jiayan Zhang, Keisuke Matsushima, Nina Kurihara, Holger Flechsig, Kenichi Umeda, Noriyuki Kodera, Kazuki Kato, Mikihiro Shibata

## Abstract

RIG-I is an innate immune receptor for viral double-stranded RNA (dsRNA). Upon binding dsRNA, RIG-I nucleates MAVS filaments through its two caspase activation and recruitment domains (2CARD). However, how the 2CARD behaves during activation has remained unclear. Using high-speed atomic force microscopy, we visualized the 2CARD in full-length RIG-I across its apo, dsRNA-bound, translocating, multimeric, and polyubiquitin-bound states. Protrusions consistent with the 2CARD appeared only transiently in the apo state, whereas a mobile 2CARD emerged upon RNA binding. During ATP-driven translocation, wild-type RIG-I moved approximately twofold slower and showed more frequent pauses and multimers than RIG-I Δ2CARD. Within multimers, 2CARDs formed a complex that polyubiquitin stabilized into larger, dsRNA-deforming assemblies. These findings reveal the 2CARD as a brake on the RIG-I motor, beyond signaling.

## Main Text

Retinoic acid–inducible gene I (RIG-I) is a cytosolic RNA sensor that initiates antiviral innate immunity (*1–3*). In the resting state, RIG-I is monomeric and adopts an autoinhibited conformation (*4*). Upon recognition of viral double-stranded RNA (dsRNA), RIG-I can assemble into short, multimeric filaments on the dsRNA in an ATP-dependent manner, bringing its N-terminal tandem caspase activation and recruitment domains (2CARD) into proximity (*5, 6*). The 2CARD modules assemble into a tetramer that is further stabilized by K63-linked polyubiquitin chains and functions as a scaffold for activating the adaptor mitochondrial antiviral-signaling protein (MAVS) (*7, 8*). MAVS filaments initiate a downstream signaling cascade involving TANK-binding kinase 1 (TBK1) and interferon regulatory factor 3 (IRF3), inducing type I interferon production and establishing an antiviral state in host cells (*1, 9, 10*).

Structural and biochemical studies have established a framework for dsRNA-induced RIG-I activation. RIG-I comprises the signaling module 2CARD, a helicase core (Hel1 and Hel2), and a C-terminal domain (CTD) (Fig. 1A) (*4, 11, 12*). In apo RIG-I, the 2CARD docks against an insertion subdomain of Hel2 (Hel2i), maintaining an autoinhibited conformation (*4*). The CTD recognizes the 5′-triphosphorylated dsRNA end, whereas the helicase core engages the dsRNA stem. Together, they encircle dsRNA in the helicase mode (*11–14*). Fluorescence resonance energy transfer (FRET) measurements showed that RNA binding is sufficient to release the 2CARD from the helicase core in the absence of ATP (*15*). However, how the exposed 2CARD moves relative to the RNA-engaged helicase core remains unclear.

**Fig. 1.**
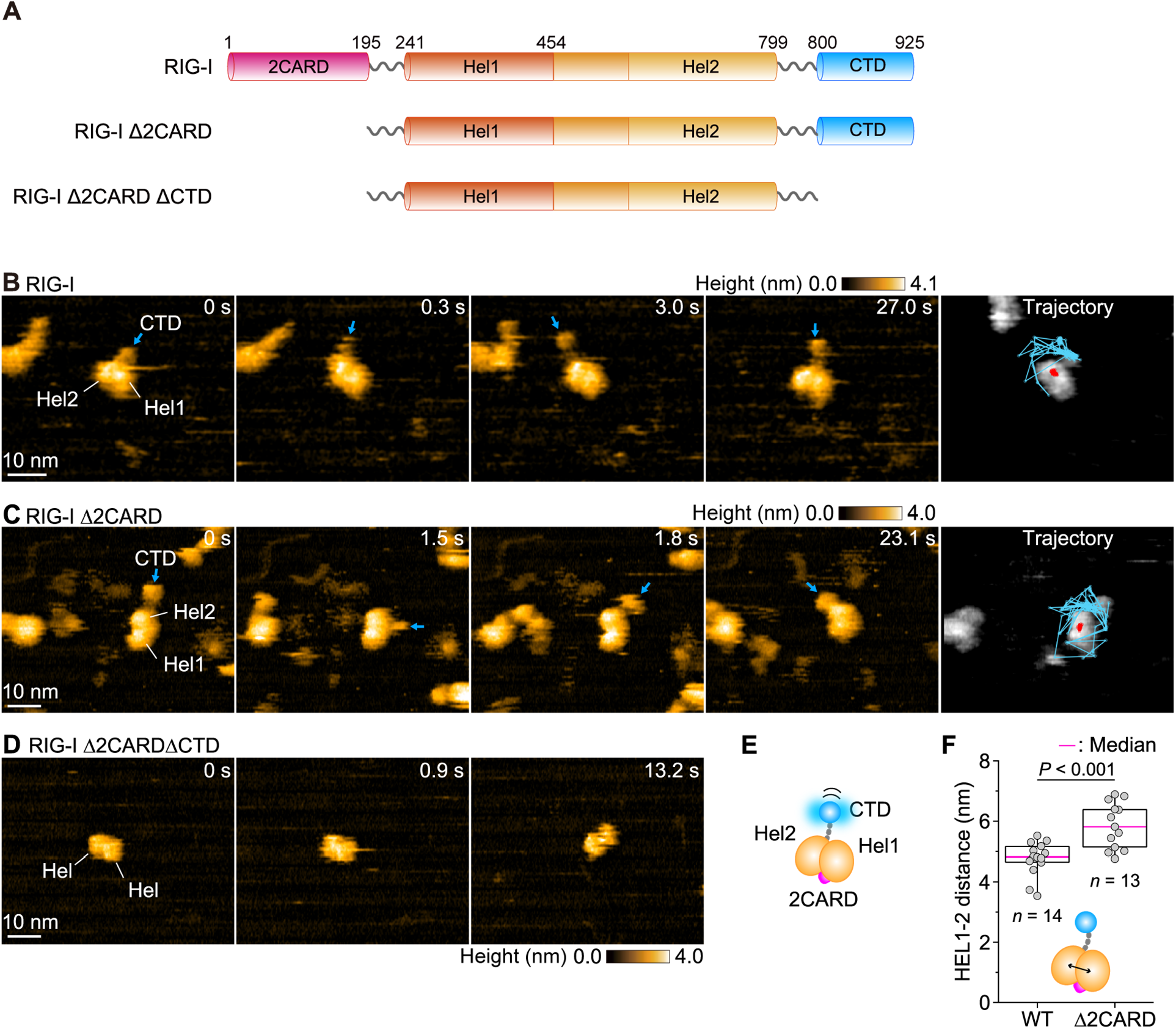
Conformational dynamics of apo RIG-I. (**A**) Domain architecture of human RIG-I and the deletion constructs used in this study. Numbers indicate residue boundaries. Hel2i (residues 465–608), an insertion within Hel2, is not resolved separately by HS-AFM and is counted within Hel2. (**B** to **D**) Sequential HS-AFM images of RIG-I (B), RIG-I Δ2CARD (C), and RIG-I Δ2CARDΔCTD (D) (movies S1, S2, and S3, respectively). Blue arrows indicate the CTD. The rightmost panels in (B) and (C) show the ∼30-s trajectories of the centroids of the CTD (blue) and helicase domains (Hel1–Hel2; red), overlaid on a representative frame. Imaging was performed at room temperature (24–26°C) in a buffer containing 20 mM HEPES-KOH (pH 7.0), 50 mM KCl, 1.5 mM MgCl_2_, 1 mM DTT, and 10% glycerol. Scan area, 60 × 48 nm^2^ in (B) and 70 × 56 nm^2^ in (C, D) (240 × 96 pixels); frame rate, 3.3 frames/s. Height scales are shown above each image series. Scale bars, 10 nm. In (D), the two helicase lobes are labeled Hel because they cannot be assigned individually in the absence of the 2CARD and CTD. (**E**) Schematic model of RIG-I domain organization. (**F**) Frame-averaged distance between Hel1 and Hel2 for RIG-I (WT) and RIG-I Δ2CARD. Each gray dot represents the mean value for one molecule (*n* = 14 for WT, 13 for Δ2CARD). In the box plot, the magenta center line represents the median, box limits delineate the 25th and 75th percentiles (IQR), and whiskers extend to the minimum and maximum values. Two-tailed Welch’s *t*-test.

A related question is how RIG-I movement is associated with its multimerization on dsRNA. RIG-I uses ATP hydrolysis to translocate along dsRNA, and the CARDs can suppress this translocation in the absence of a 5′-triphosphate end (*16*). Biochemical and kinetic studies further linked translocation, throttling at the 5′-triphosphate end, and multimerization on dsRNA (*5, 6, 14*). However, how the exposed 2CARD affects movement along dsRNA and how this movement relates to contacts between neighboring RIG-I molecules remain unclear.

Here, we used high-speed atomic force microscopy (HS-AFM) (*17–20*) to examine full-length RIG-I in its apo, dsRNA-bound, translocating, multimeric, and polyubiquitin-bound states. We observed a mobile CTD and transient protrusions consistent with 2CARD exposure in apo RIG-I, and a mobile 2CARD in the RNA-bound helicase mode. Compared with RIG-I lacking the 2CARD (RIG-I Δ2CARD), wild-type RIG-I translocated more slowly, spent more time in pauses, and was more frequently observed in multimers. Neighboring 2CARDs formed transient contacts that were stabilized by polyubiquitin, which also supported larger 2CARD clusters. These observations link the signaling module of RIG-I to its movement and organization on dsRNA.

## Results

### The CTD is mobile, whereas the 2CARD is rarely resolved in apo RIG-I

We purified full-length human RIG-I (fig. S1A) and imaged single molecules of apo RIG-I by HS-AFM. Each molecule showed two adjacent elliptical domains, from which a spherical domain protruded and fluctuated (Fig. 1B; movie S1). To assign these domains, we imaged two deletion constructs. RIG-I Δ2CARD retained both the elliptical domains and the mobile spherical domain (Fig. 1C; movie S2), whereas RIG-I Δ2CARDΔCTD showed only the two elliptical domains (Fig. 1D; movie S3). These comparisons assigned the spherical domain to the CTD and the two elliptical domains to Hel1 and Hel2. The domain connected to the CTD by a linker was assigned as Hel2 (Fig. 1E). The CTD–helicase core distance was 8.1 ± 1.4 nm, and the radius of gyration (*R*g) of the CTD trajectory was 3.6 ± 0.6 nm (fig. S1, B and C), indicating substantial CTD mobility relative to the helicase core.

Comparison of RIG-I and RIG-I Δ2CARD revealed no persistent protrusion attributable to the 2CARD (Fig. 1, B and C). Occasionally, however, a protrusion consistent with the 2CARD became transiently visible in full-length RIG-I (fig. S1D; movie S1, example 3). These observations are consistent with transient exposure of the 2CARD in the apo state (*21*). Finally, the frame-averaged Hel1–Hel2 distance was shorter in RIG-I (4.8 ± 0.57 nm; *n* = 14 molecules) than in RIG-I Δ2CARD (5.9 ± 0.76 nm; *n* = 13 molecules) (Fig. 1F), suggesting that the 2CARD contributes to a more compact arrangement of the helicase domains in the apo state.

### The 2CARD is exposed in the RNA-bound helicase mode

To examine the domain organization of RNA-bound RIG-I, we imaged RIG-I with a 312-bp dsRNA bearing 5′-triphosphate (5′ppp-dsRNA) in the absence of ATP. RIG-I bound 5′ppp-dsRNA more frequently than dsRNA with a 5′-hydroxyl end (5′OH-dsRNA) (fig. S2, A and B), and most bound molecules were located at the dsRNA end (Fig. 2A). In one configuration, the spherical CTD capped the dsRNA end and fluctuated around it (Fig. 2B; movie S4, examples 1 and 2). In a second configuration, RIG-I appeared as an elliptical particle at the dsRNA end, accompanied by a mobile spherical domain (Fig. 2C; movie S4, examples 3 and 4, magenta arrow). RIG-I Δ2CARD formed a similar elliptical particle but lacked the mobile spherical domain (Fig. 2D; movie S5), supporting assignment of this domain to the 2CARD. We then generated simulated AFM images (*22, 23*) from a cryo-electron microscopy (cryo-EM) structure of full-length RIG-I bound to dsRNA (PDB 8SD0) (*24*). These matched the elliptical particle (cross-correlation 0.82; Fig. 2E), supporting its assignment to the helicase mode, in which Hel1, Hel2, and the CTD have wrapped around the RNA duplex (*11–13, 25*). Both the CTD-capped configuration and the helicase mode were observed in the absence of ATP. A mobile 2CARD was visible in the helicase mode but was not resolved as a separate domain in the CTD-capped configuration (Fig. 2, B and C). These observations are consistent with previous ensemble FRET measurements indicating RNA-induced release of the 2CARD from the helicase core (*15*). Our images provide a direct view of the released 2CARD in RNA-bound RIG-I, although the transition from the CTD-capped configuration to the helicase mode was not directly observed.

**Fig. 2.**
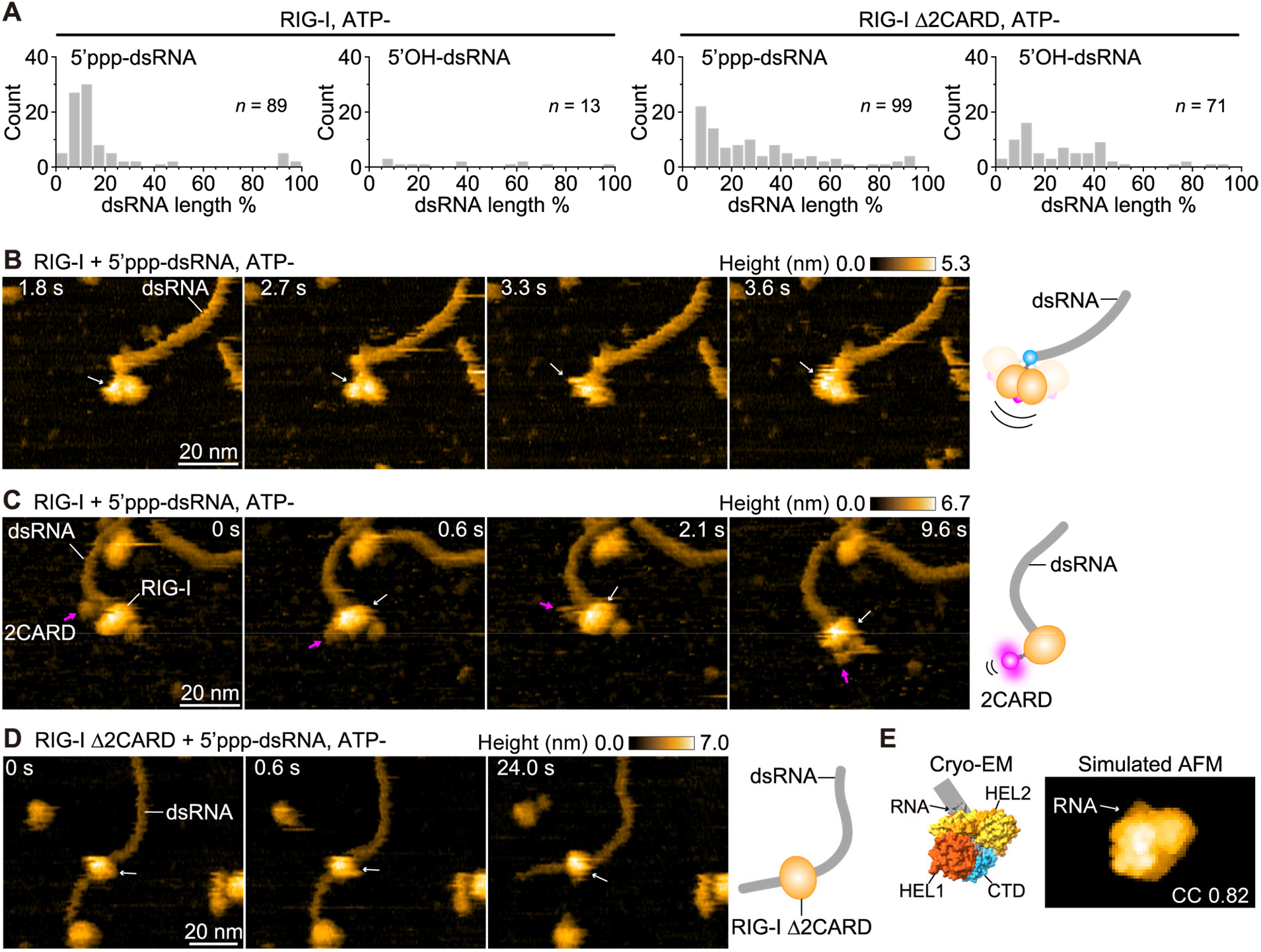
Dynamics of single RIG-I molecules on dsRNA in the absence of ATP. (**A**) Histograms showing the binding position of RIG-I along 5′ppp-dsRNA or 5′OH-dsRNA, expressed as percent dsRNA length from one end, for RIG-I (left) and RIG-I Δ2CARD (right). Numbers of molecules analyzed (*n*) are indicated in each panel. (**B** and **C**) Sequential HS-AFM images of RIG-I (white arrow) bound to 5′ppp-dsRNA. RIG-I associates with the dsRNA end via the CTD and fluctuates continuously (B; movie S4, examples 1 and 2), or wraps around the dsRNA with the 2CARD (magenta arrow) accompanying the helicase core (C; movie S4, examples 3 and 4). Right, schematic model (also in D). (**D**) Sequential HS-AFM images of RIG-I Δ2CARD (white arrow) bound to 5′ppp-dsRNA (movie S5). (**E**) Cryo-EM structure of full-length RIG-I bound to RNA, in which the 2CARD is not resolved (PDB 8SD0) (*24*), and the corresponding simulated AFM image. Cross-correlation (CC) between the simulated and experimental AFM images, 0.82. Imaging was performed at room temperature (24–26°C) in a buffer containing 20 mM HEPES-KOH (pH 7.0), 150 mM KCl, 1.5 mM MgCl_2_, 1 mM DTT, and 10% glycerol. Scan area, 80 × 64 nm^2^ in (B, C) and 100 × 80 nm^2^ in (D) (240 × 96 pixels); frame rate, 3.3 frames/s. Height scales are shown above each image series. Scale bars, 20 nm.

Whereas wild-type RIG-I bound predominantly at the end of 5′ppp-dsRNA, RIG-I Δ2CARD was also observed at internal regions of 5′ppp-dsRNA and bound 5′OH-dsRNA (Fig. 2A; fig. S2, C and D). Together with the wider Hel1–Hel2 separation in apo RIG-I Δ2CARD (Fig. 1F), these findings suggest that the 2CARD contributes to both helicase-domain organization and preferential binding to the dsRNA end under these conditions. Whether the wider Hel1–Hel2 separation observed in the apo state causes the altered RNA-binding pattern remains to be determined.

### The 2CARD slows RIG-I translocation and increases pausing

To examine RIG-I translocation and the associated 2CARD dynamics, we imaged RIG-I on 5′ppp-dsRNA in the presence of ATP. RIG-I was observed both at the dsRNA end and at internal regions (Fig. 3A; fig. S3A). Most molecules adopted the helicase mode with an exposed 2CARD. The 2CARD moved parallel to the dsRNA, at a distance of 9.9 ± 1.5 nm from the helicase core, with an *R*g of 4.2 ± 0.9 nm (Fig. 3, B and C).

**Fig. 3.**
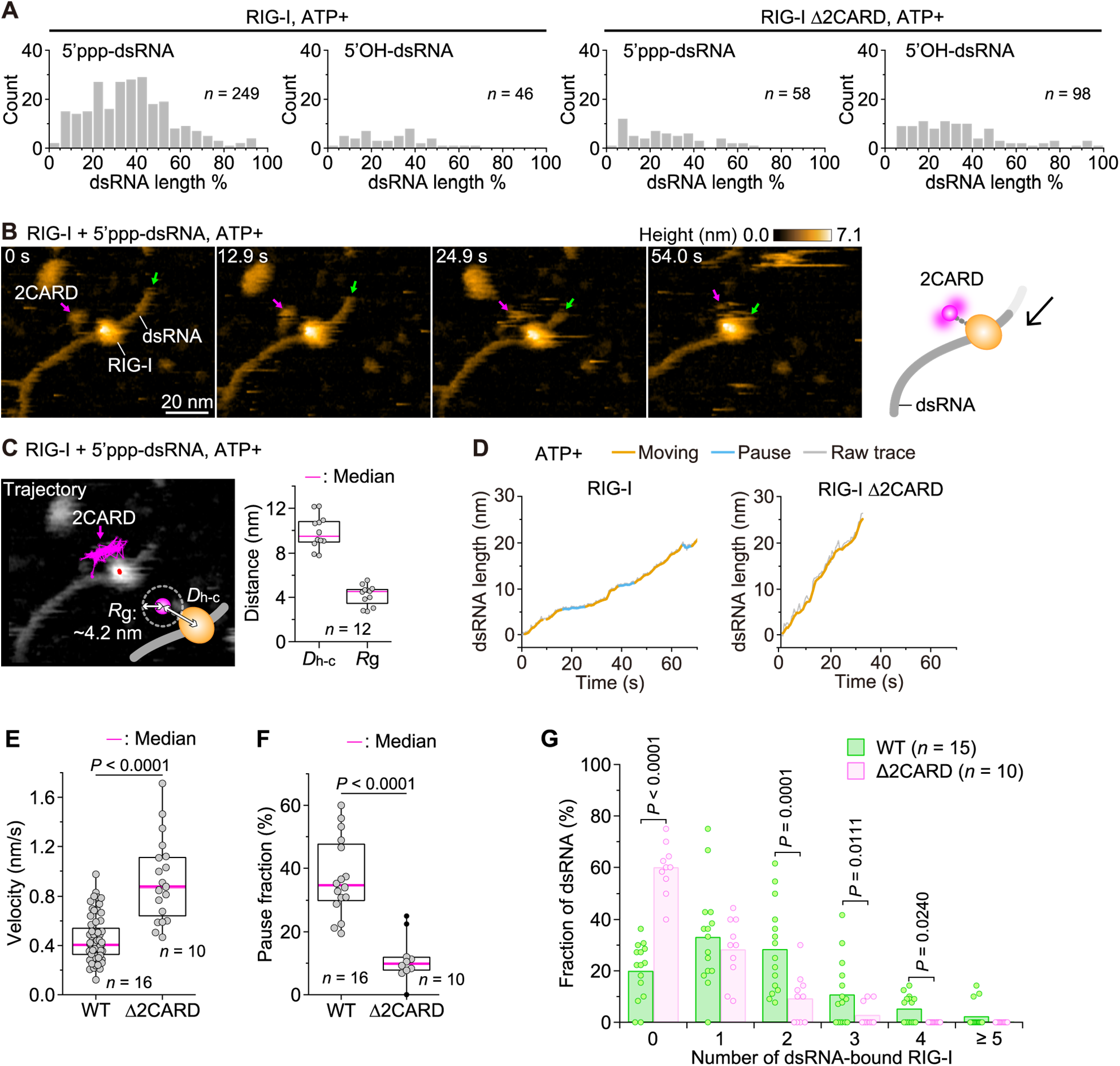
Dynamics of single RIG-I molecules on dsRNA in the presence of ATP. (**A**) Histograms showing the binding position of RIG-I along 5′ppp-dsRNA or 5′OH-dsRNA, expressed as percent dsRNA length from one end, for RIG-I (left) and RIG-I Δ2CARD (right). (**B**) Sequential HS-AFM images showing translocation of 5′ppp-dsRNA (end marked by green arrow) driven by a single RIG-I molecule (2CARD, magenta arrow; movie S6). Scan area, 80 × 64 nm^2^ (240 × 96 pixels); frame rate, 3.3 frames/s. The height scale is shown above the image series. Scale bars, 20 nm. Right, schematic model. (**C**) Left, ∼30-s trajectories of the 2CARD (magenta) and the helicase core centroid (red), overlaid on a representative frame. *D*_h-c_, distance between the 2CARD and helicase core centroids; *R*_g_, radius of gyration of the 2CARD trajectory. Right, box plot of *D*_h-c_ and *R*_g_. (**D**) A representative time course of the length of 5’ppp-dsRNA translocated by a single RIG-I (left) and RIG-I Δ2CARD (right) molecule. Orange, moving segments; light blue, pauses; gray, raw trace. Traces for other analyzed molecules are shown in fig. S3, B and D. (**E**) Translocation velocity of 5′ppp-dsRNA driven by RIG-I (WT) and RIG-I Δ2CARD. Two-tailed Welch’s *t*-test. (**F**) Fraction of time spent in pauses during translocation of 5’ppp-dsRNA driven by RIG-I (WT) and RIG-I Δ2CARD. Two-tailed Welch’s *t*-test. (**G**) Distribution of the number of RIG-I molecules bound per dsRNA molecule (six categories, 0 to ≥5) for RIG-I (WT) (*n* = 15 independent experiments; 162 dsRNA molecules in total) and RIG-I Δ2CARD (*n* = 10 independent experiments; 100 dsRNA molecules in total). Wilcoxon test with Benjamini-Hochberg correction. In box plots (C, E, F), the magenta center line represents the median, box limits delineate the 25th and 75th percentiles (IQR), whiskers extend to the minimum and maximum values, and black dots represent outliers identified by the Tukey criterion (beyond 1.5× the IQR from the box). In (C) and (F), each gray dot represents one molecule; in (E), each gray dot represents one velocity measurement, with multiple measurements obtained per molecule. The difference in velocity remained significant when tested on per-molecule mean velocities (16 and 10 molecules for the wild type and Δ2CARD, respectively; Welch’s t-test, *P* = 0.0011). In (G), each dot represents one independent experiment. The numbers of molecules analyzed are indicated in each panel (C, F). Imaging was performed at room temperature (24–26°C) in a buffer containing 20 mM HEPES-KOH (pH 7.0), 150 mM KCl, 1.5 mM MgCl_2_, 1 mM DTT, 10% glycerol, and 0.2 mM ATP.

Successive HS-AFM images captured unidirectional movement of the dsRNA relative to RIG-I (Fig. 3B; movie S6). RIG-I was immobilized on the AFM substrate, whereas the dsRNA remained free to move; thus, the observed RNA movement reported relative translocation in this surface-bound assay. Tracing the dsRNA contour length from RIG-I to the end over time yielded trajectories for individual molecules (Fig. 3D; additional traces in fig. S3B). The translocation velocity increased with ATP concentration (fig. S3C), supporting ATP-dependent movement. RIG-I Δ2CARD also supported unidirectional translocation (Fig. 3D; fig. S3, D and E; movie S7), indicating that the 2CARD is dispensable for movement under these conditions.

However, wild-type RIG-I translocated more slowly (0.39 ± 0.18 nm/s) than RIG-I Δ2CARD (0.81 ± 0.22 nm/s) (Fig. 3E), even though wild-type RIG-I hydrolyzed ATP at a slightly higher rate than RIG-I Δ2CARD in a separate biochemical assay (fig. S3F). The dsRNA repeatedly stopped and restarted, and the fraction of observation time classified as pauses was higher for wild-type RIG-I (37 ± 12%) than for RIG-I Δ2CARD (11 ± 7%) (Fig. 3F). For wild-type RIG-I, pauses lasted 7.2 ± 4.1 s on average and occurred at a frequency of 0.091 ± 0.088 events/s (fig. S3, G and H). The pause fraction did not increase monotonically with ATP concentration over the range examined (fig. S3I). Together, these observations indicate that the 2CARD slows translocation and increases the time spent in pauses.

We next examined whether these differences were associated with multimerization by counting the number of RIG-I molecules bound to individual dsRNA molecules (Fig. 3G). The fractions of dsRNA molecules bound by two, three, or four RIG-I molecules were higher for the wild type (28 ± 17%, 11 ± 13%, and 5 ± 5%, respectively) than for RIG-I Δ2CARD (9 ± 10%, 3 ± 5%, and 0%, respectively). Conversely, 88% of dsRNA molecules had no more than one RIG-I Δ2CARD molecule bound, compared with 53% for the wild type. Thus, slower translocation and more time spent in pauses in wild-type RIG-I were associated with a greater prevalence of multimers on dsRNA.

### Neighboring 2CARDs associate in pairs within RIG-I multimers

Since 2CARD tetramers serve as scaffolds for activating MAVS signaling (*7, 8*), we next examined the organization and dynamics of the 2CARDs within RIG-I multimers on dsRNA (Fig. 4; movies S8 to S11). When two RIG-I molecules were bound next to each other on dsRNA, their 2CARDs repeatedly associated and separated (Fig. 4A; movie S8), indicating transient contacts under the imaging conditions. In assemblies of three RIG-I molecules, two 2CARDs formed a complex while the third remained separate (Fig. 4B; movie S9). In the larger assemblies shown, containing four, five, or eight RIG-I molecules, the 2CARDs were distributed on both sides of the dsRNA, and the resolved 2CARD complexes comprised pairs (Fig. 4, C to E; movies S10 and S11). These observations suggest that neighboring 2CARDs form transient pairs within the imaged multimers (Fig. 4F).

**Fig. 4.**
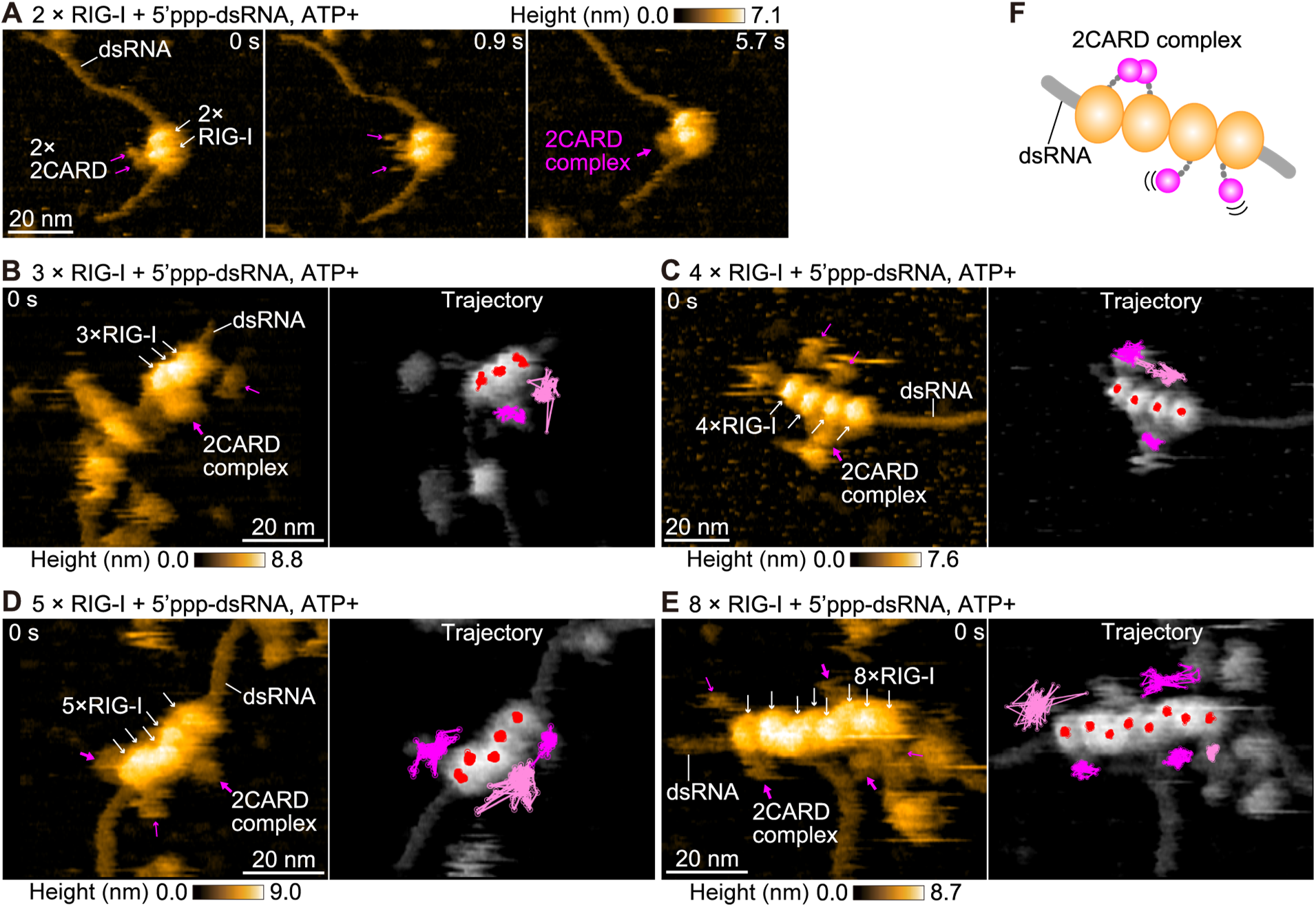
Dynamics of the 2CARD in RIG-I multimers assembled on dsRNA. (**A**) Sequential HS-AFM images of two RIG-I molecules (white arrows) assembled on 5′ppp-dsRNA. The two 2CARD modules (magenta arrows) repeatedly associated with and dissociated from one another (movie S8). (**B** to **E**) Left, HS-AFM images of three (B), four (C), five (D), and eight (E) RIG-I molecules (white arrows) assembled on 5′ppp-dsRNA, with the 2CARD complex indicated (magenta arrow). Right, ∼30-s trajectories of the individual RIG-I core centroids (red), the single 2CARD (light magenta), and the complex formed by two 2CARD modules (dark magenta) (B, movie S9; C, movie S10; D and E, movie S11, examples 3 and 4, respectively). (**F**) Schematic model showing the arrangement of 2CARD within a RIG-I multimer assembled on dsRNA. Imaging was performed at room temperature (24–26°C) in a buffer containing 20 mM HEPES-KOH (pH 7.0), 150 mM KCl, 1.5 mM MgCl_2_, 1 mM DTT, 10% glycerol, and 0.1–2.0 mM ATP. Scan area, 80 × 64 nm^2^ (240 × 96 pixels) except for (C), 100 × 80 nm^2^; frame rate, 3.3 frames/s. Height scales are shown above or below each image series. Scale bars, 20 nm.

### Polyubiquitin stabilizes the 2CARD complex and expands it beyond two molecules

K63-linked polyubiquitin has been shown to support RIG-I activation and stabilize 2CARD tetramers that nucleate MAVS filaments (*7, 8, 26*). To examine its effects on 2CARD dynamics, we imaged RIG-I multimers on 5′ppp-dsRNA in the presence of polyubiquitin (Fig. 5A; movie S12). Tracking adjacent 2CARDs showed more persistent contacts in the presence of polyubiquitin (Fig. 5B). The fraction of imaging time classified as dissociated decreased from 19 ± 4% without polyubiquitin to 0.76 ± 0.10% with polyubiquitin (Fig. 5C), indicating that polyubiquitin stabilized contacts between 2CARDs under these conditions.

**Fig. 5.**
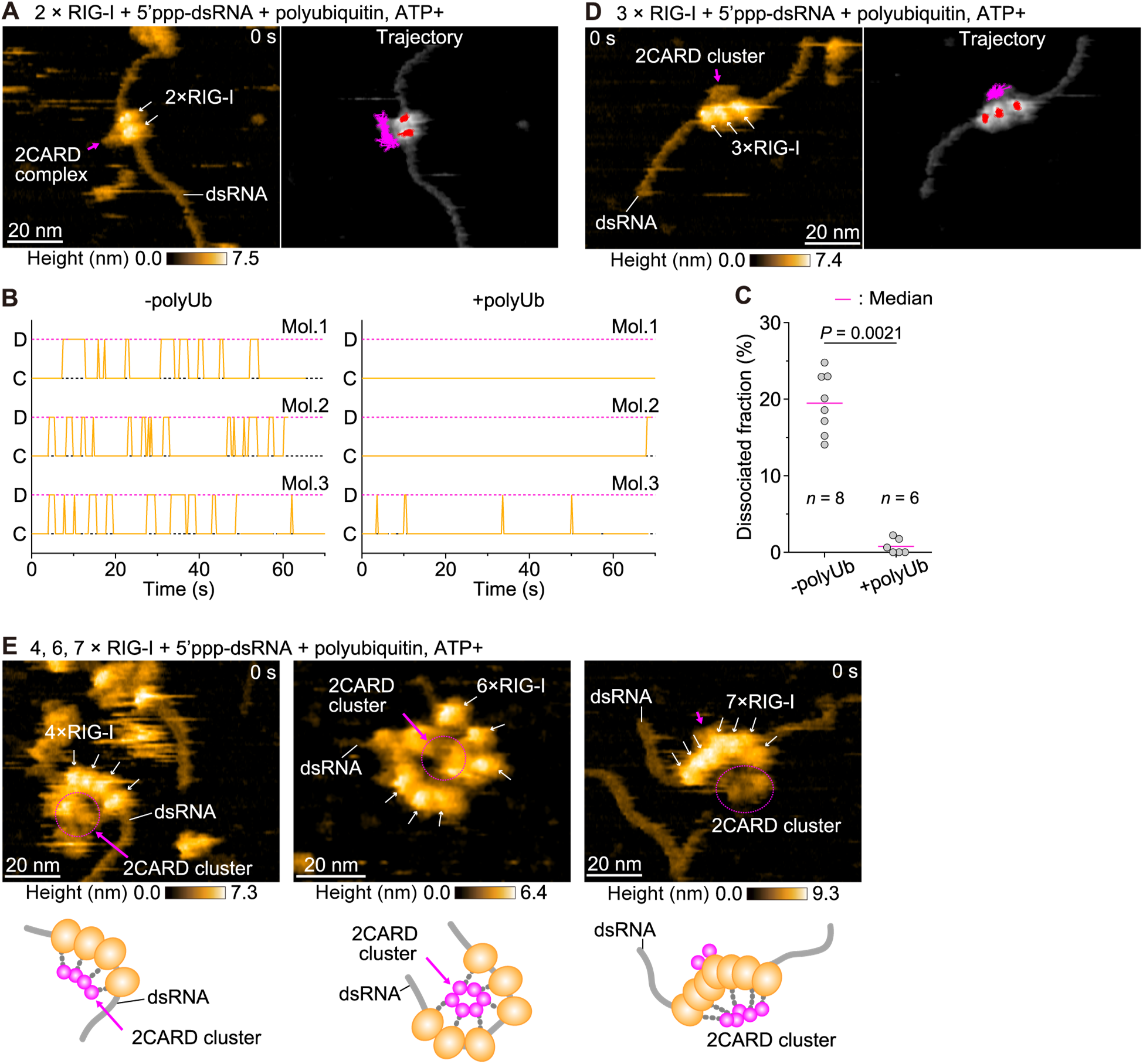
Effect of polyubiquitin on the dynamics of the 2CARD in multimeric RIG-I assembled on dsRNA. (**A**) HS-AFM images of two RIG-I molecules (white arrows) assembled on 5’ppp-dsRNA in the presence of polyubiquitin, with the associated 2CARD complex (magenta arrow). Trajectory panel shows ∼30-s trajectories of the individual RIG-I core centroids (red) and the 2CARD complex (magenta) (movie S12). (**B**) Representative traces of 2CARD complex dynamics over time for individual pairs of neighboring RIG-I molecules in the absence (−polyUb, left) or presence (+polyUb, right) of polyubiquitin. D, dissociated; C, in contact. (**C**) Fraction of imaging time during which the 2CARD complex was dissociated, in the absence (−polyUb) or presence (+polyUb) of polyubiquitin. Two-tailed Welch’s t-test. (**D**) HS-AFM image of three RIG-I molecules (white arrows) assembled on 5’ppp-dsRNA in the presence of polyubiquitin, with the associated 2CARD cluster (magenta arrow). Trajectory panel shows ∼30-s trajectories of the individual RIG-I core centroids (red) and the 2CARD cluster (magenta) (movie S13). (**E**) HS-AFM images of four, six, or seven RIG-I molecules (white arrows) assembled on 5’ppp-dsRNA in the presence of polyubiquitin, with the associated 2CARD cluster (magenta arrow/circle). Schematic models illustrate the corresponding assemblies. Movies are shown for the four- and seven-molecule assemblies (movie S14). Imaging was performed at room temperature (24–26°C) in a buffer containing 20 mM HEPES-KOH (pH 7.0), 150 mM KCl, 1.5 mM MgCl_2_, 1 mM DTT, 10% glycerol, and 0.2 mM ATP, supplemented with polyubiquitin where indicated. Scan area, 100 × 80 nm² (240 × 96 pixels) except for (E, middle), 80 × 64 nm^2^; frame rate, 3.3 frames/s. Height scales are shown below each image series. Scale bars, 20 nm.

Higher-order 2CARD assemblies were also observed in the presence of polyubiquitin. In assemblies of three RIG-I molecules on the same dsRNA, all three 2CARDs formed a single cluster in *cis* in the presence of polyubiquitin (Fig. 5D; movie S13), whereas only two associated with each other in its absence (Fig. 4B). In higher-order RIG-I assemblies, the 2CARDs formed a common cluster associated with curved or looped dsRNA (Fig. 5E; movie S14). These assemblies also included RIG-I molecules whose helicase cores were not visibly bound to dsRNA. In addition, 2CARDs from RIG-I molecules bound to separate dsRNAs associated in *trans*, cross-bridging the two RNAs (fig. S4 and movie S12, example 3). Together, these observations indicate that RIG-I 2CARDs can oligomerize in multiple configurations in the presence of polyubiquitin, both in *cis* on the same dsRNA and in *trans* between separate dsRNAs.

## Discussion

Using HS-AFM, we addressed two open questions in RIG-I activation: how the 2CARD behaves as RIG-I engages dsRNA, and how translocating RIG-I molecules assemble into multimers on dsRNA. In apo RIG-I, protrusions consistent with the 2CARD were only transiently visible, whereas the 2CARD was mobile in the RNA-bound helicase mode. Wild-type RIG-I translocated more slowly and spent more time in pauses than RIG-I Δ2CARD, and multimers were more prevalent with the wild type. Within these multimers, neighboring 2CARDs formed transient contacts that were stabilized by polyubiquitin. Together, these observations support a role for the 2CARD in regulating RIG-I motion and molecular organization in addition to its signaling function.

In resting RIG-I, the 2CARD docks against Hel2i to maintain an autoinhibited conformation (*4*). Deletion of the 2CARD increased the Hel1–Hel2 separation and broadened RNA binding to include internal regions of 5′ppp-dsRNA and 5′OH-dsRNA (Figs. 1F and 2A). These observations suggest that 2CARD docking favors CTD-mediated recognition of 5′ppp ends while limiting direct engagement of dsRNA stems by the helicase core. The mobility of the CTD observed here likely facilitates recognition of 5′ppp-dsRNA ends. This gating would prevent inappropriate binding to self-RNAs, including dephosphorylated and capped RNAs, and thereby serve as a checkpoint for RNA recognition (*27*). Upon RNA recognition, RIG-I releases the 2CARD from the helicase core, making it available for intermolecular association. This model is consistent with previous FRET measurements and our observation of a mobile 2CARD in RNA-bound RIG-I (*15*). Together, these observations indicate that the 2CARD contributes to discrimination of viral from self-RNA and becomes exposed for intermolecular association upon viral RNA recognition.

The 2CARD also influenced RIG-I behavior after RNA recognition. Previous studies established CARD-dependent regulation of translocation and ATPase-driven RNA proofreading (*14, 16*). In our experiments, wild-type RIG-I moved more slowly and spent more time paused than RIG-I Δ2CARD, despite showing slightly higher ATPase activity in a separate assay (fig. S3F). This raises the possibility that ATP hydrolysis and translocation are partially uncoupled in wild-type RIG-I, although the different assay conditions preclude a direct comparison of ATP consumption per unit displacement. LGP2, a related RNA sensor that lacks the 2CARD, showed apparently continuous, unidirectional translocation in previous HS-AFM observations (*20*), supporting a role for the 2CARD in RIG-I pausing. A paused RIG-I molecule can act as a transient roadblock that increases encounters with molecules translocating behind it, thereby favoring multimer formation on dsRNA. This mechanism parallels the recently described LGP2-mediated control of MDA5, in which LGP2 has been proposed to block MDA5 movement and promote filament assembly (*28*).

Because we could not observe the assembly process directly, we built a mathematical model in which RIG-I molecules translocate along a 106-nm dsRNA and multimerize upon contact (fig. S5A, Materials and Methods). This model reproduced the overall trend of the dsRNA-bound RIG-I distribution observed by HS-AFM (fig. S5B). In the model, raising the velocity to the Δ2CARD value while keeping the wild-type pauses did not prevent multimerization (fig. S5C). This indicated that pausing rather than slow movement drives assembly. In contrast, removing the pauses at the wild-type velocity abolished multimerization entirely (fig. S5D). Pauses were therefore required for contact-dependent association, whereas the reduced velocity had a smaller effect. Together, these observations and simulations support a model in which 2CARD-dependent pausing promotes RIG-I assembly by increasing opportunities for intermolecular contact on dsRNA.

CARD–CARD interactions between the exposed 2CARDs provide a potential link between RNA-bound assembly and downstream signaling. K63-linked polyubiquitin stabilized transient 2CARD contacts and supported higher-order clusters, with interactions occurring both in *cis* on the same dsRNA and in *trans* between different dsRNAs. These versatile binding modes may allow RIG-I to organize signaling assemblies on RNAs of different lengths, from short duplexes to longer dsRNAs. Considering that RIG-I 2CARD tetramers are sufficient to induce MAVS filament formation (*7, 8*), the polyubiquitin-stabilized higher-order 2CARD oligomers observed here are likely to activate MAVS signaling.

Several features of the imaging assay constrain the interpretation. We did not directly observe the transition from CTD-mediated end recognition to the helicase mode or the recruitment of additional molecules to a paused RIG-I. Surface immobilization may affect movement and domain accessibility; RNA motion relative to immobilized RIG-I therefore need not reproduce its kinetics in solution. A substrate that immobilizes dsRNA while allowing RIG-I to move and bind from solution could help resolve recruitment dynamics. In addition, imaging does not measure ATP turnover, and the pause fraction depends on trajectory analysis and classification. These limitations should be addressed when testing the proposed link between 2CARD-dependent motion and multimerization.

Following individual RIG-I molecules in real time revealed transient, asynchronous behaviors that ensemble and structural measurements cannot resolve. Together, these observations reveal the dynamic behavior of the multifunctional RIG-I 2CARD and support a model in which it acts as a checkpoint for self–non-self RNA discrimination, a brake on translocation that favors multimer formation, and a scaffold for MAVS signaling (fig. S6).

## Acknowledgments

Claude Opus 4.8 (Anthropic) was used to assist with language editing and proofreading of the manuscript. The authors reviewed and verified all content and take full responsibility for it. No generative AI was used for data generation, analysis, or interpretation. We thank Prof. Toshio Ando at Kanazawa University for providing HS-AFM apparatus, and Yui Okada and Youko Mikami at Kanazawa University for collecting and analyzing HS-AFM data. This work was supported by the WISE Program of Kanazawa University by MEXT.

## Funding

The World Premier International Research Center Initiative (WPI), MEXT, Japan (MS)

JSPS KAKENHI grant numbers JP24K21942 (MS)

JSPS KAKENHI grant numbers JP25H00972 (MS)

JSPS KAKENHI grant numbers JP26K23141 (MS)

JSPS KAKENHI grant numbers JP25H01315 (KK)

Takeda Science Foundation (KK, MS)

JST SPRING JPMJSP2135 (SK, KM)

JST ERATO JPMJER2403 (MS)

JST FOREST JPMJFR240V (KK)

## Author contributions

Conceptualization: MS

Data curation: SK, JZ, MS

Formal analysis: SK, KM, MS

Funding acquisition: KK, MS

Investigation: SK, JZ, NK, MS

Methodology: SK, KM, HF, KU, NK, KK, MS

Project administration: KK, MS

Resources: KK, MS

Software: SK, KU, MS

Supervision: KK, MS

Validation: SK, MS

Visualization: SK, KK, MS

Writing – original draft: MS

Writing – review & editing: KK, MS

## Competing interests

The authors declare that they have no competing interests.

## Data, code, and materials availability

All data are available in the main text or the supplementary materials. The simulation code is available at Zenodo (*29*).

## Supplementary Materials

Materials and Methods

Figs. S1 to S6

References (30–36)

Movies S1 to S14

## Notes

### Competing Interest Statement

The authors have declared no competing interest.

