## Supplemental file for "Dual role of the RIG-I 2CARD as a signaling module and a brake on dsRNA translocation"

**This PDF file includes:**

Materials and Methods
Figs. S1 to S6
References (30–36)

**Other Supplementary Materials for this manuscript include the following:**
Movies S1 to S14

### Materials and Methods

Plasmids

The coding sequence of human RIG-I (DDX58) was cloned into the pE-SUMO expression vector to generate an N-terminal His-SUMO-tagged RIG-I construct. RIG-I Δ2CARD (residues 204–925) and RIG-I Δ2CARDΔCTD (residues 204–793) constructs were generated by deletion of the corresponding 2CARD and CTD regions from the full-length RIG-I sequence. Plasmids were constructed by PCR-based cloning and verified by Sanger sequencing.

Purification of RIG-I

RIG-I WT, RIG-I Δ2CARD, and RIG-I Δ2CARDΔCTD proteins were expressed in Rosetta2 (DE3) in LB medium (Nacalai Tesque). Cells were grown at 37°C to an optical density at 600 nm (OD600) of 0.6 and induced with 0.25 mM isopropyl β-D-1-thiogalactopyranoside (IPTG; Nacalai Tesque) at 18°C overnight. Cells were lysed by sonication in buffer A (20 mM Tris-HCl, pH 8.0, 20 mM imidazole, 300 mM NaCl, 10% glycerol, and 3 mM 2-mercaptoethanol) supplemented with 2 mM $MgCl_2$, and the lysate was clarified by centrifugation at 40,000 × g for 30 min at 4°C. The supernatant was applied to Ni-NTA Agarose resin (QIAGEN), and bound protein was eluted with buffer A supplemented with 300 mM imidazole. The SUMO tag was cleaved by incubation with Ulp1 protease at 4°C overnight. The eluted fraction was diluted with buffer B (10 mM sodium phosphate, pH 7.5, 0.5 mM EDTA, and 3 mM 2-mercaptoethanol) to adjust the final NaCl concentration to 150 mM and loaded onto a HiTrap Heparin column (Cytiva) pre-equilibrated with buffer C (10 mM sodium phosphate, pH 7.5, 0.5 mM EDTA, 150 mM NaCl, and 3 mM 2-mercaptoethanol). Bound protein was eluted with a linear NaCl gradient from 150 mM to 2 M NaCl using buffer B and buffer D (10 mM sodium phosphate, pH 7.5, 0.5 mM EDTA, 2 M NaCl, and 3 mM 2-mercaptoethanol). Peak fractions containing the target proteins were concentrated and further purified by size-exclusion chromatography on a Superdex 200 Increase 10/300 GL column (Cytiva) equilibrated with buffer E (20 mM HEPES, pH 7.5, 150 mM NaCl, and 2 mM dithiothreitol).

RNA preparation

312 bp dsRNAs used in this study were prepared by in vitro T7 transcription. The templates for RNA synthesis were generated by PCR amplification. The sequence of the 312-bp dsRNA was taken from the first 300 bp of the human MDA5 gene, flanked by 5′-gggaga and 5′-tctccc. The two complementary RNA strands were co-transcribed, and the

duplex was purified using 6% acrylamide gel electrophoresis. RNA was gel-extracted using the Model 422 Electro-Eluter electroelution kit (Biorad), ethanol precipitated, and stored at −80°C until use. For 5′OH-dsRNA, the purified 312 bp dsRNA was treated with Calf intestinal alkaline phosphatase (CIP, NEB) to remove 5′-triphosphate moiety, and was re-purified using QIAquick Nucleotide removal kit (QIAGEN).

In vitro ATPase Assay

ATPase activity of RIG-I and RIG-I Δ2CARD proteins was measured using a phosphate-release assay with BIOMOL Green reagent (Enzo Life Sciences). RIG-I or RIG-I Δ2CARD protein was diluted to 400 nM in ATPase reaction buffer containing 20 mM HEPES (pH 7.5), 150 mM NaCl, 1.5 mM $MgCl_2$, 2 mM DTT, and 0.4 mM ATP. The 312-bp dsRNA was diluted to 2 ng/μL in the same reaction buffer without ATP. For each reaction, protein solution was mixed with either RNA solution or RNA-free buffer. The final concentrations were 200 nM protein, 1 ng/μL RNA, and 0.2 mM ATP. Reactions were incubated at 37°C for 5 min and subsequently transferred to a clear-bottom 96-well plate. BIOMOL Green reagent was added to each reaction, followed by incubation at room temperature for 20 min. Absorbance was measured at 620 nm. ATP-containing reaction buffer without protein was used for background correction. Each condition was measured with four technical replicates ($n = 4$).

HS-AFM Equipment

HS-AFM imaging was performed on a custom-built microscope (*17, 30*), as described previously (*19, 20, 31*). Imaging was performed in tapping mode using MNOIC AC7 cantilevers (length 6–7 μm, width 2 μm, thickness 100 nm). Cantilever oscillation was monitored by an optical-lever system: an infrared laser (780 nm, 0.8 mW) was focused onto the back of the cantilever through a 60× objective (CFI S Plan Fluor ELWD 60X, Nikon), and the reflected beam was directed onto a PIN photodiode. The cantilevers had a spring constant of ~100 pN/nm and, in liquid, a resonance frequency of 400–600 kHz and a quality factor of ~2. An additional tip (~300 nm long) was grown at the cantilever apex by electron-beam deposition on a field-emission scanning electron microscope (Verios 5UC, Thermo Fisher Scientific) and sharpened to an apex radius of ~1 nm by plasma etching (Tergeo, PIE Scientific); all images were acquired with this tip. The free-oscillation amplitude was kept below 1 nm, and the set-point amplitude was ~90% of the free amplitude. HS-AFM operation and data acquisition were controlled with UMEX. To minimize tip-induced damage, images were acquired only during the forward (trace) scan, with the tip retracted from the sample during the backward (retrace) scan (only-trace imaging mode) (*32*).

HS-AFM Observations

*Apo RIG-I*

Apo RIG-I WT, RIG-I Δ2CARD, and RIG-I Δ2CARDΔCTD were observed on freshly cleaved bare mica. A 3-μL aliquot of protein at 20–25 nM was deposited on the mica and allowed to adsorb for 5 min at room temperature. Unbound molecules were washed away with 40 μL of observation buffer A (20 mM HEPES-KOH, pH 7.0, 50 mM KCl, 1.5 mM $MgCl_2$, 1 mM DTT, 10% glycerol), and imaging was performed in the same buffer.

*RIG-I–dsRNA Complex*

RIG-I–dsRNA complexes were observed on mica modified with 3-aminopropyltriethoxysilane (APTES; AP-mica). APTES (0.0033% in Milli-Q water) was applied to freshly cleaved mica for 3 min at room temperature, and the surface was washed with 40 μL of Milli-Q water followed by 40 μL of observation buffer. AP-mica was prepared freshly and used immediately.

Complexes were formed with 5′-triphosphate (5′ppp) dsRNA or 5′-OH dsRNA. In the absence of ATP, RIG-I and dsRNA were mixed at a molar ratio of 20:1 or 30:1 in observation buffer B (20 mM HEPES-KOH, pH 7.0, 150 mM KCl, 1.5 mM $MgCl_2$, 1 mM DTT, 10% glycerol) and incubated for 5 min at room temperature. A 3-μL aliquot was applied to AP-mica for 5 min, unbound molecules were washed away with 40 μL of observation buffer B, and imaging was performed in the same buffer. In the presence of ATP, RIG-I and dsRNA were mixed at a 30:1 molar ratio in observation buffer B supplemented with 0.1–2.0 mM ATP and incubated for 10 min at room temperature. A 3-μL aliquot was applied to AP-mica for 5 min, unbound molecules were washed away with 40 μL of the same buffer, and imaging was performed in that buffer. For the ATP-titration experiments (fig. S3, C and I), in which ATP was raised up to 1.5 mM, the $MgCl_2$ concentration was increased to 10 mM to maintain sufficient free $Mg^{2+}$, because ATP chelates $Mg^{2+}$; all other experiments used 1.5 mM $MgCl_2$.

*In the Presence of Polyubiquitin*

RIG-I, dsRNA, and 0.2 mM ATP were mixed as under the ATP-containing condition (RIG-I:dsRNA = 30:1) and incubated for 10 min at room temperature. Polyubiquitin was then added to 140 nM, 280 nM, or 1.6 μM, and the mixture was incubated for a further 5 min. A 3-μL aliquot was applied to AP-mica for 5 min, and unbound molecules were washed away with 40 μL of observation buffer B containing 0.2 mM ATP. Final concentrations were 60–120 nM RIG-I and ~3 nM dsRNA.

All HS-AFM imaging was performed at 24–26°C. At least three independent experiments were performed for each condition, and representative images and movies are shown.

HS-AFM Image Processing and Data Analysis

HS-AFM images were processed and analyzed using Fiji/ImageJ (NIH) (*33*), R, Python, and custom scripts. Background was subtracted for each image with the Subtract Background function in Fiji/ImageJ (rolling-ball radius, 200 pixels), and lateral drift between successive frames was corrected with the Template Matching and Slice Alignment plugin (*34*). For apo RIG-I, the coordinates of the globular structures corresponding to Hel1, Hel2, and the CTD, and the center of the 2CARD, were tracked frame-by-frame in a semi-automated manner with the MTrackJ plugin (*35*). Quantitative analyses, including inter-domain distances within RIG-I and dsRNA velocity, were restricted to movies and images with clearly identifiable molecular shapes and no marked tip artifacts, instrumental drift, or molecular overlap.

*Analysis of CTD and 2CARD trajectories*

To quantify the trajectories of the CTD and the 2CARD, we computed the radius of gyration, $R_g$, as the root-mean-square displacement from the average position:

$\langle\vec{R}\rangle$, i.e., $R_g^2 = (1/N)\sum_{n=1}^{N}\left(\vec{R}_k(n) - \langle\vec{R}\rangle\right)^2$,

where $\langle\vec{R}\rangle$ is the average position of the CTD or the 2CARD. $R_g$ was calculated from HS-AFM movies at 3.3 frames/s over ~70–100 frames (about 30 s).

*RIG-I Binding Position*

For each condition (RIG-I or RIG-I Δ2CARD with 5′ppp-dsRNA or 5′OH-dsRNA), the total length of dsRNA in each HS-AFM image was measured and normalized to 0–100%. A height profile was obtained along the dsRNA contour, and positions with a height ≥4.5 nm above the substrate were defined as RIG-I binding sites. Because the two dsRNA ends could not be distinguished in HS-AFM images, the position of the first RIG-I molecule was measured from the nearer end. Repeating the analysis with thresholds of 5.0 and 5.5

nm gave consistent results. Adjacent peaks arising from the same RIG-I molecule were merged on the basis of the apparent width of RIG-I (~7.0 nm).

*ATP-Dependent Analysis of dsRNA Translocation*

To quantify the translocation velocity and pausing of RIG-I on dsRNA, the dsRNA contour length from one end to the center of the RIG-I molecule was measured in each frame and traced over time. First, a 3-point moving average was applied to reduce noise. A line was then fitted to the distance–time relationship within a ~1-s window centered on each time point, and the absolute value of the slope defined the local velocity (nm/s) at that point. The distribution of local velocities was bimodal, and a two-component Gaussian mixture model (GMM) (*36*) separated it into a low-velocity (pause) and a high-velocity (moving) component. The boundary between the two components was 0.158 nm/s: velocities at or below this value were classified as pauses, and those above it as moving. The velocity reported in Fig. 3E, which was also used as the moving velocity in the simulation, is the mean of the local velocities classified as moving. Low-velocity intervals shorter than 3 s were excluded from both the velocity and the pause analyses; only pauses lasting 3 s or longer were included as pauses. Because the dsRNA moved unidirectionally relative to RIG-I, the absolute value of the local slope was used.

*Analysis of 2CARD Complex Dynamics*

To assess 2CARD complex formation within RIG-I multimers, the 2CARDs of adjacent molecules were classified as dissociated or in contact. A height profile was obtained across the 2CARD protrusions: two clearly separated peaks defined the dissociated state, and a single or continuous peak defined the in-contact state. Frame-by-frame classification was used to follow the dynamics of 2CARD dissociation and complex formation over time.

Simulation of RIG-I Multimerization on dsRNA

To examine the correlation between the pauses and multimerization of RIG-I on dsRNA observed by HS-AFM, we performed a JavaScript-based simulation. Each frame corresponded to 0.1 s, and simulations were run for up to 400 s. Based on the HS-AFM data, the dsRNA was modeled as a 106-nm straight line, and RIG-I as a circle 7.0 nm in diameter. In the simulation, RIG-I bound only to one end of the dsRNA and moved unidirectionally toward the other end. RIG-I molecules were introduced sequentially: each new molecule bound to the end 20–50 s (random) after the previous one. This interval was chosen to allow the preceding RIG-I molecule to move away from the end before the next one binds, preventing multimerization at the dsRNA end and allowing it to be examined in the central region of the dsRNA. The maximum number of molecules introduced per run was 20, but given the introduction interval (mean 35 s) and the observation limit (400 s), on average about 11 molecules were actually introduced per run (fig. S5A). Molecules that reached the far end of the dsRNA were considered to have dissociated. Translocation used the mean velocity of each construct as a constant value [WT, 0.39 nm/s; Δ2CARD, 0.81 nm/s (Fig. 3E)]. At each molecule's introduction, its pause duration (7.2 ± 4.1 s; fig. S3G) and frequency (0.091 ± 0.088 events/s; fig. S3H) were each sampled once at random from the measured values. When the center-to-center distance between two RIG-I molecules on the dsRNA became ≤7.0 nm, the molecules were defined as bound and recorded as a complex. After binding, each molecule continued to move independently according to its own velocity and pause parameters. We refer to a group of molecules connected by such binding as a "complex" (a single unbound molecule is counted as a complex of size 1). To prevent molecules from overlapping, the step size of a trailing molecule was adjusted. This upper limit on the trailing molecule's step size was recalculated at each step from its position relative to the molecule ahead, so that the trailing molecule could advance again automatically once the molecule ahead moved and opened a gap. When the calculated step size exceeded this limit, movement was truncated at the limit.

We analyzed the simulated multimerization of RIG-I on the dsRNA (fig. S5A) as follows. One 400-s simulation constituted a trial. Trials were grouped into blocks, and each block was plotted as one data point (*n* = 20 blocks in every analysis). Because the two analyses extract different quantities from each trial, the number of trials per block differed, as described below. For the multimer distribution (fig. S5B), we recorded the multimerization state of RIG-I on the dsRNA every 10 s from 40 s to 400 s of each trial, giving 37 records per trial. We excluded the first 30 s because few molecules had entered the dsRNA during this initial loading period. The records from 15 trials were combined and categorized into five classes (1, 2, 3, 4, or ≥5 molecules); one block therefore comprised 15 trials, matching the 15 independent HS-AFM experiments. For each block, the fraction of records in each class was calculated (Fraction of dsRNA, %), and twenty blocks were plotted (fig. S5B; *n* = 20).

To assess the effects of velocity and pausing, we recorded, for each trial, the largest multimer size that any RIG-I reached, which gives a single value per trial. Because each trial contributes only one value, 200 trials were combined into one block to obtain a stable fraction, and twenty blocks were plotted (fig. S5, C and D; $n = 20$). To test velocity, we changed the velocity to the Δ2CARD value while keeping the other parameters (introduction interval, pause frequency, and pause duration) at the wild-type values (fig. S5C); to test pausing, we set both the pause frequency and duration to zero (fig. S5D). In each case, the wild-type condition was simulated in parallel using the same trials and the same random draws.

Quantification and Statistical Analysis

Statistical calculations were performed in R, Python, OriginPro 2026b (OriginLab Corporation) and Igor Pro 10 (WaveMetrics, USA), as appropriate. All tests were two-sided, with a significance level of $\alpha = 0.05$. For multiple comparisons, corrected rather than raw $P$ values are reported. Where a test assumed normality or homogeneity of variance, these were confirmed with the Shapiro–Wilk and Levene's tests, respectively. The statistical test, correction method, sample size, and data units for each analysis are given in the corresponding figure legend. Unless otherwise stated, values are reported as mean ± SD.

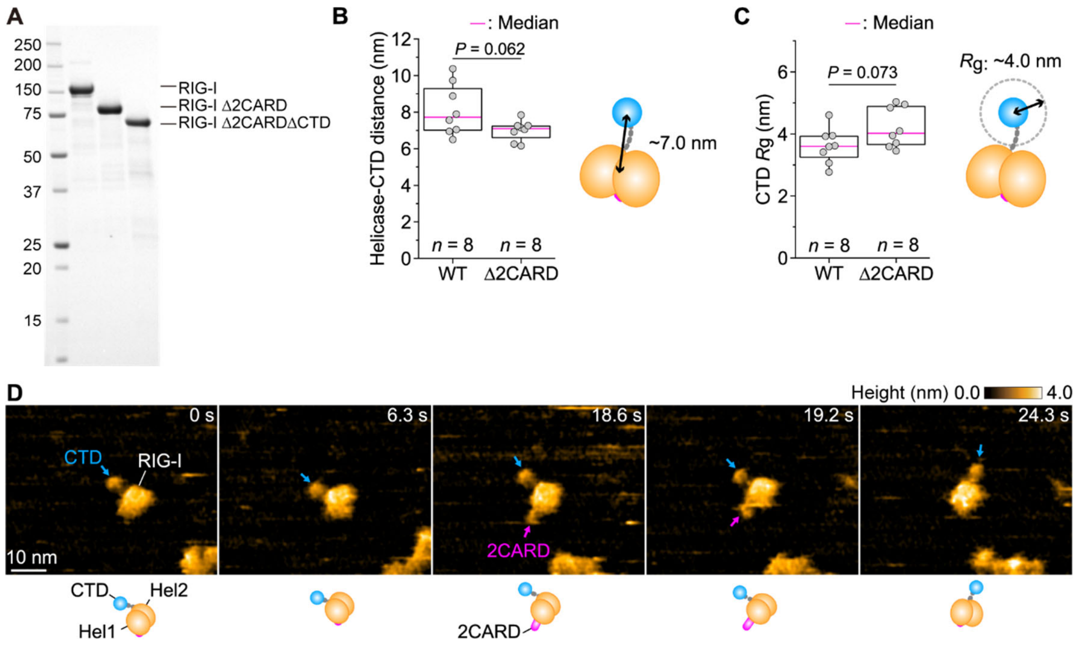


**Fig. S1. Sample preparation and dynamics of apo RIG-I.**

(**A**) Purity of RIG-I, RIG-I Δ2CARD, and RIG-I Δ2CARDΔCTD assessed by SDS-PAGE with Coomassie Brilliant Blue staining.

(**B**) Distance between the helicase core and the CTD for RIG-I (WT) and RIG-I Δ2CARD. Two-tailed Welch's *t*-test.

(**C**) Radius of gyration (*R*g) of the CTD trajectory for RIG-I (WT) and RIG-I Δ2CARD, calculated from ~70 consecutive frames per molecule. Two-tailed Welch's *t*-test.

(**D**) Sequential HS-AFM images of RIG-I (movie S1, third molecule), showing exposure of the 2CARD (magenta arrow) as the CTD (blue arrow) moves away from the helicase core. Bottom, corresponding schematic models.

Numbers of molecules analyzed (*n*) are indicated in each panel. In box plots (B, C), the magenta center line represents the median, box limits delineate the 25th and 75th percentiles (IQR), whiskers extend to the minimum and maximum values, and gray dots represent individual molecules. Imaging was performed at room temperature (24–26°C) in a buffer containing 20 mM HEPES-KOH (pH 7.0), 50 mM KCl, 1.5 mM $MgCl_2$, 1 mM DTT, and 10% glycerol. Height scales are shown above each image series. Scale bars, 10 nm.

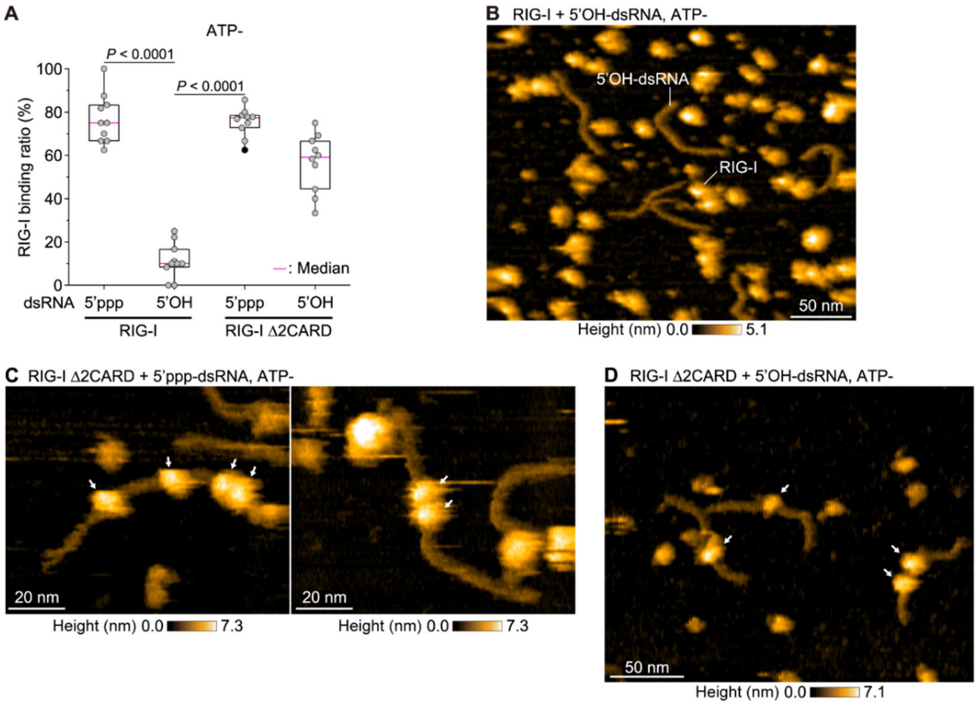


**Fig. S2. HS-AFM control experiments, and binding of RIG-I Δ2CARD at internal dsRNA positions.**

(**A**) Fraction of dsRNA molecules bound by RIG-I or RIG-I Δ2CARD, for 5′ppp-dsRNA and 5′OH-dsRNA, determined from HS-AFM images acquired over different scan areas across three independent experiments ($n$ = 10 dsRNA molecules per condition). In box plots, the magenta center line represents the median, box limits delineate the 25th and 75th percentiles (IQR), whiskers extend to the minimum and maximum values, and gray dots represent individual dsRNA molecules. Kruskal-Wallis test with Dunn's post hoc test.

(**B**) HS-AFM image of RIG-I incubated with 5′OH-dsRNA, showing lower occupancy of RIG-I on 5′OH-dsRNA under these conditions.

(**C**) HS-AFM images showing multiple RIG-I Δ2CARD molecules bound to 5′ppp-dsRNA, including two molecules bound in close proximity within an internal region (movie S5, third example).

(**D**) HS-AFM image of RIG-I Δ2CARD bound to 5′OH-dsRNA, showing binding at an internal region of the dsRNA.

Imaging was performed at room temperature (24–26°C) in a buffer containing 20 mM HEPES-KOH (pH 7.0), 150 mM KCl, 1.5 mM $MgCl_2$, 1 mM DTT, and 10% glycerol. Scan area, 300 × 240 nm$^2$ (280 × 112 pixels) in (B), 100 × 80 nm$^2$ (240 × 96 pixels) in (C), and 250 × 200 nm$^2$ in (D) (240 × 96 pixels); frame rate, 2.0 frames/s in (B) and (D), 3.3 frames/s in (C). Height scales are shown below each image series. Scale bars, 50 nm in (B, D) and 20 nm in (C).

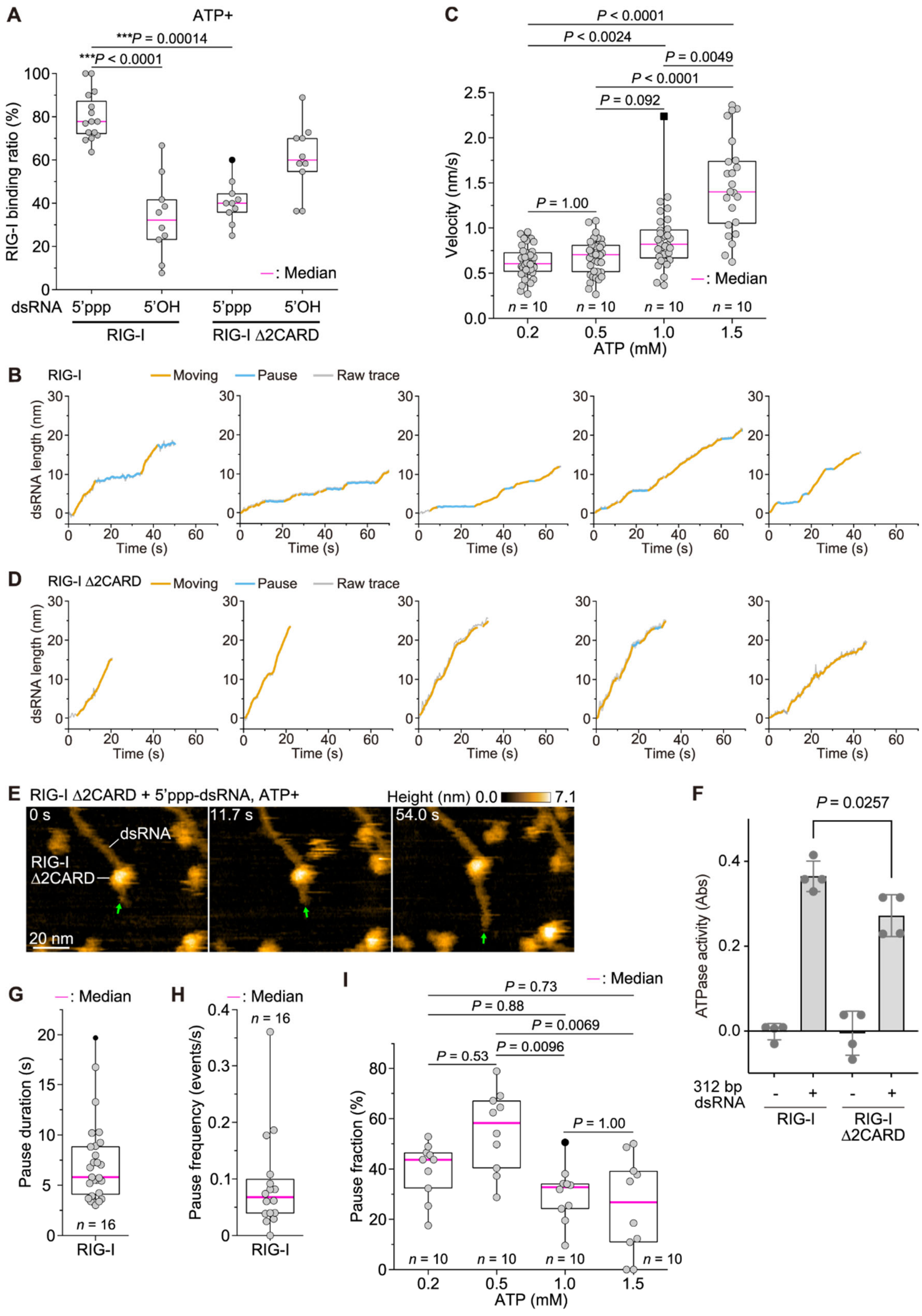
A
ATP+
***P = 0.00014
***P < 0.0001
RIG-I binding ratio (%)
—: Median
dsRNA 5'ppp 5'OH 5'ppp 5'OH
RIG-I
RIG-I Δ2CARD
C
P < 0.0001
P < 0.0024
P = 0.0049
P < 0.0001
P = 0.092
P = 1.00
Velocity (nm/s)
—: Median
n = 10
ATP (mM)
B
RIG-I
Moving
Pause
Raw trace
dsRNA length (nm)
Time (s)
D
RIG-I Δ2CARD
Moving
Pause
Raw trace
dsRNA length (nm)
Time (s)
E
RIG-I Δ2CARD + 5'ppp-dsRNA, ATP+
Height (nm) 0.0 7.1
0 s
11.7 s
54.0 s
dsRNA
RIG-I
Δ2CARD
20 nm
F
P = 0.0257
ATPase activity (Abs)
312 bp
dsRNA
RIG-I
RIG-I
Δ2CARD
G
—: Median
Pause duration (s)
n = 16
RIG-I
H
—: Median
n = 16
Pause frequency (events/s)
RIG-I
I
—: Median
P = 0.73
P = 0.88
P = 0.0069
P = 0.0096
P = 0.53
P = 1.00
Pause fraction (%)
n = 10
ATP (mM)

**Fig. S3. Translocation dynamics and ATPase activity of RIG-I and RIG-I Δ2CARD.**
(**A**) Fraction of dsRNA molecules bound by RIG-I or RIG-I Δ2CARD, for 5′ppp-dsRNA and 5′OH-dsRNA in the presence of ATP, determined from HS-AFM images acquired over different scan areas across three independent experiments ($n$ = 10 dsRNA molecules per condition). Kruskal-Wallis test with Dunn's post hoc test.
(**B** and **D**) Five representative time courses of the length of 5'ppp-dsRNA translocated by single RIG-I (B) and RIG-I Δ2CARD (D) molecules in the presence of 0.2 mM ATP. Orange, moving segments; light blue, pauses; gray, raw trace.
(**C**) Translocation velocity of 5′ppp-dsRNA driven by RIG-I (WT) as a function of ATP concentration, measured in 10 mM $MgCl_2$. Kruskal-Wallis test with Dunn's post hoc test and Bonferroni correction.
(**E**) Sequential HS-AFM images of RIG-I Δ2CARD translocating along 5′ppp-dsRNA in the presence of 1 mM ATP (end marked by green arrow; movie S7). Scan area, 100 × 80 $nm^2$ (240 × 96 pixels); frame rate, 3.3 frames/s. The height scale is shown above each image series. Scale bars, 20 nm.
(**F**) ATPase activity of RIG-I and RIG-I Δ2CARD, measured with a malachite-green (BIOMOL Green) phosphate-release assay in the presence of 5′ppp-dsRNA. Wild-type RIG-I hydrolyzed ATP at a slightly higher rate than RIG-I Δ2CARD. Two-tailed Welch's $t$-test. Data are mean ± SD of four technical replicates ($n$ = 4).
(**G**) Duration of individual pauses during translocation of 5′ppp-dsRNA by RIG-I (WT) at 0.2 mM ATP.
(**H**) Frequency of pauses during translocation of 5′ppp-dsRNA by RIG-I (WT) at 0.2 mM ATP.
(**I**) Fraction of time spent in pauses during translocation of 5'ppp-dsRNA by RIG-I (WT) as a function of ATP concentration, measured in 10 mM $MgCl_2$. Kruskal-Wallis test with Dunn's post hoc test and Bonferroni correction.
In box plots (A, C, G, H, I), the magenta center line represents the median, box limits delineate the 25th and 75th percentiles (IQR), whiskers extend to the minimum and maximum values, and black dots represent outliers identified by the Tukey criterion (beyond 1.5× the IQR from the box). Gray dots represent individual dsRNA molecules in (A), individual velocity measurements in (C), individual pauses in (G), and individual RIG-I molecules in (H) and (I). Numbers of molecules analyzed ($n$) are indicated in each panel. Imaging was performed at room temperature (24–26°C) in a buffer containing 20 mM HEPES-KOH (pH 7.0), 150 mM KCl, 1.5 mM $MgCl_2$ unless otherwise noted, 1 mM DTT, and 10% glycerol, supplemented with ATP as indicated in each panel.

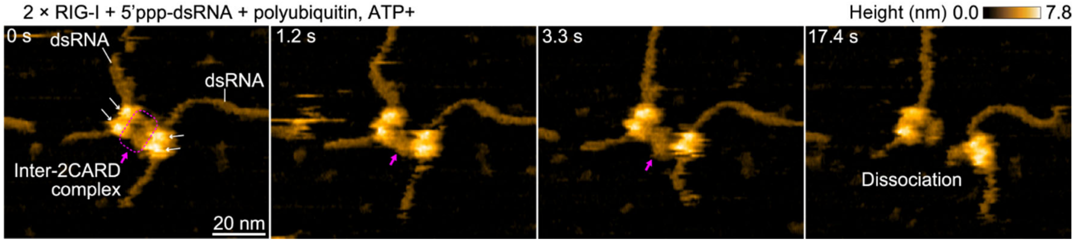


**Fig. S4. An intermolecular 2CARD complex forms between RIG-I molecules on separate dsRNAs.**

Sequential HS-AFM images of two RIG-I molecules (white arrows), each bound to a different 5′ppp-dsRNA, in the presence of polyubiquitin and ATP. Their 2CARDs associated in *trans*, forming a complex (magenta arrow and dashed outline) that bridged the two dsRNAs and later dissociated (movie S12, example 3). Imaging was performed at room temperature (24–26°C) in a buffer containing 20 mM HEPES-KOH (pH 7.0), 150 mM KCl, 1.5 mM $MgCl_2$, 1 mM DTT, 10% glycerol, 0.2 mM ATP, and polyubiquitin. Scan area, 100 × 80 $nm^2$ (240 × 96 pixels); frame rate, 3.3 frames/s. The height scale is shown above the image series. Scale bar, 20 nm.

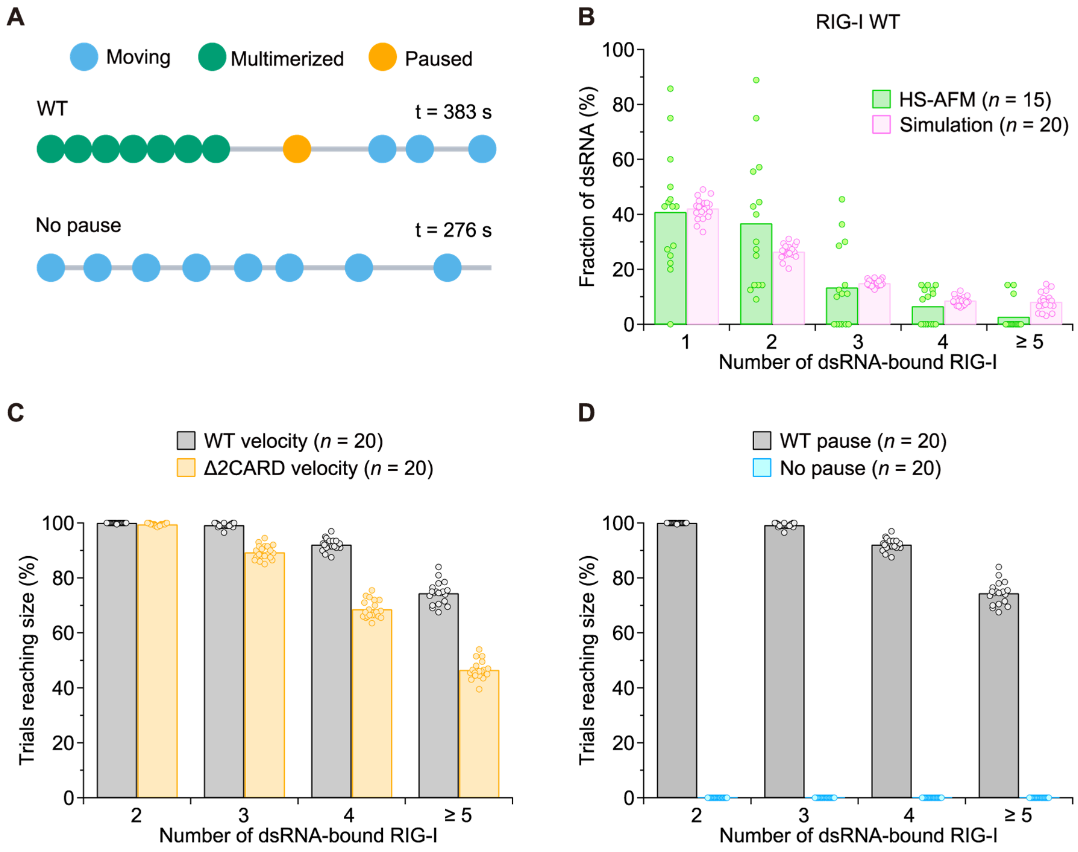


**Fig. S5. Simulation of RIG-I multimerization on dsRNA.**

(**A**) Representative snapshots of simulations under the wild-type and no-pause conditions, showing RIG-I molecules that are moving, paused, or multimerized on a single dsRNA. The simulation time is indicated.

(**B**) Distribution of the number of dsRNA-bound RIG-I molecules at each time point, for RIG-I WT, compared between HS-AFM ($n$ = 15 independent experiments; 162 dsRNA molecules in total) and simulation ($n$ = 20 blocks). Each point is one experiment or one block.

(**C**) Fraction of simulation trials in which RIG-I reached each multimer size, comparing the wild-type velocity with the Δ2CARD velocity ($n$ = 20 blocks each).

(**D**) Fraction of simulation trials in which RIG-I reached each multimer size, comparing the wild-type pauses with the no-pause condition ($n$ = 20 blocks each).

In (**C**) and (**D**), all parameters except the one tested were held at the wild-type values. Bars show the mean; each point is one block.

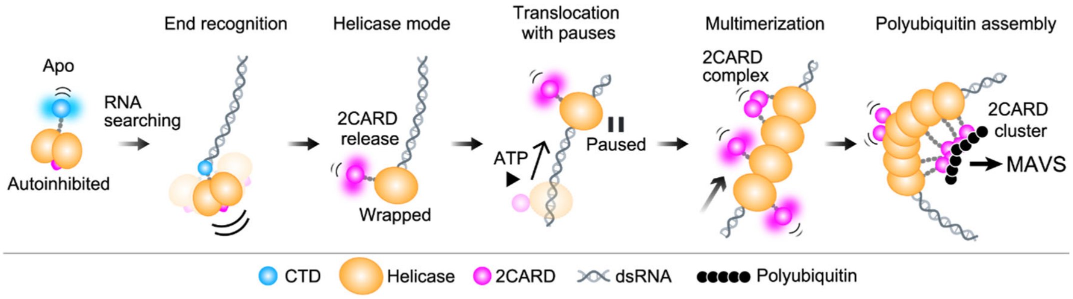


**Fig. S6. Model for 2CARD dynamics during RIG-I activation.**
In apo RIG-I, the helicase domain holds the 2CARD in an autoinhibited conformation and keeps Hel1 and Hel2 compact, which may contribute to preferential binding at the dsRNA end. Recognition of the 5′-triphosphate end by the CTD and formation of the helicase mode release the 2CARD as a mobile module. RIG-I translocates along the dsRNA using ATP hydrolysis, and the 2CARD brakes this motion and introduces pauses. During these pauses, following RIG-I molecules assemble into a multimer on the dsRNA, and their 2CARDs form a complex. Polyubiquitin stabilizes and enlarges this complex into a cluster large enough to deform the dsRNA. Such clusters may provide a platform for MAVS filament nucleation; this activity was not tested here.

**Supplementary Movie legends**

**Movie S1.** HS-AFM movies of three representative molecules of apo RIG-I on bare mica. Blue arrows indicate the CTD. Scan area, 60 × 48 $nm^2$ (molecules 1, 3) or 70 × 56 $nm^2$ (molecule 2); image size, 240 × 96 pixels (molecules 1, 3), 120 × 96 pixels (molecule 2) as acquired during HS-AFM observations; frame rate, 3.3 frames/s.

**Movie S2.** HS-AFM movies of three representative molecules of apo RIG-I Δ2CARD on bare mica. Blue arrows indicate the CTD. Scan area, 70 × 56 $nm^2$; image size, 240 × 96 pixels as acquired during HS-AFM observations; frame rate, 3.3 frames/s.

**Movie S3**. HS-AFM movies of three representative molecules of apo RIG-I Δ2CARDΔCTD on bare mica. Scan area, 70 × 56 $nm^2$; image size, 240 × 96 pixels as acquired during HS-AFM observations; frame rate, 3.3 frames/s.

**Movie S4.** HS-AFM movies of four representative molecules of RIG-I bound to 5'ppp-dsRNA in the absence of ATP on AP-mica. White and magenta arrows indicate RIG-I and 2CARD, respectively. Scan area, 80 × 64 $nm^2$; image size, 240 × 96 pixels as acquired during HS-AFM observations; frame rate, 3.3 frames/s.

**Movie S5.** HS-AFM movies of three representative molecules of RIG-I Δ2CARD bound to 5′ppp-dsRNA in the absence of ATP on AP-mica. In molecule 3, two RIG-I Δ2CARD molecules are bound in close proximity at the center of the dsRNA. Scan area, 100 × 80 $nm^2$; image size, 240 × 96 pixels as acquired during HS-AFM observations; frame rate, 3.3 frames/s.

**Movie S6.** HS-AFM movies of three representative molecules of RIG-I bound to 5′ppp-dsRNA in the presence of ATP on AP-mica (0.2 mM ATP for molecules 1 and 2; 2 mM ATP for molecule 3). Green arrows indicate the end of the dsRNA. Scan area, 80 × 64 $nm^2$; image size, 240 × 96 pixels as acquired during HS-AFM observations; frame rate, 3.3 frames/s.

**Movie S7.** HS-AFM movies of three representative molecules of RIG-I Δ2CARD bound to 5′ppp-dsRNA in the presence of ATP on AP-mica (0.2 mM ATP for molecule 1; 1 mM ATP for molecules 2 and 3). Green arrows indicate the end of the dsRNA. Scan area, 100 × 80 $nm^2$; image size, 240 × 96 pixels as acquired during HS-AFM observations; frame rate, 2 frames/s for molecule 1 and 3.3 frames/s for molecules 2 and 3.

**Movie S8.** HS-AFM movies of three representative examples of two RIG-I molecules bound to 5′ppp-dsRNA in the presence of ATP on AP-mica (1 mM ATP for examples 1 and 3; 2 mM ATP for example 2). Magenta arrows indicate the 2CARD complex. Scan area, 80 × 64 $nm^2$; image size, 240 × 96 pixels as acquired during HS-AFM observations; frame rate, 3.3 frames/s.

**Movie S9.** HS-AFM movies of three representative examples of three RIG-I molecules bound to 5′ppp-dsRNA in the presence of ATP on AP-mica (0.2 mM ATP for examples 1 and 2; 0.1 mM ATP for example 3). Magenta arrows indicate the 2CARD complex and an uncomplexed 2CARD. Scan area, 80 × 64 $nm^2$ for examples 1 and 2, and 100 × 80 $nm^2$ for example 3; image size, 240 × 96 pixels as acquired during HS-AFM observations; frame rate, 3.3 frames/s.

**Movie S10.** HS-AFM movies of three representative examples of four RIG-I molecules bound to 5′ppp-dsRNA in the presence of ATP on AP-mica (0.1 mM ATP for example 1; 0.2 mM ATP for examples 2 and 3). Magenta arrows indicate the 2CARD complex and an uncomplexed 2CARD. Scan area, 100 × 80 $nm^2$ for examples 1 and 3, and 80 × 64 $nm^2$ for example 2; image size, 240 × 96 pixels as acquired during HS-AFM observations; frame rate, 3.3 frames/s.

**Movie S11.** HS-AFM movies of four representative examples of five RIG-I molecules (examples 1 and 2) and eight RIG-I molecules (examples 3 and 4) bound to 5′ppp-dsRNA in the presence of 0.2 mM ATP on AP-mica. Magenta arrows indicate the 2CARD complex and an uncomplexed 2CARD. Scan area, 80 × 64 $nm^2$ for examples 1–3 and 100 × 80 $nm^2$ for example 4; image size, 240 × 96 pixels as acquired during HS-AFM observations; frame rate, 3.3 frames/s.

**Movie S12.** HS-AFM movies of three representative examples of two RIG-I molecules bound to 5′ppp-dsRNA in the presence of 0.2 mM ATP and polyubiquitin on AP-mica. Magenta arrows indicate the 2CARD complex. Scan area, 100 × 80 $nm^2$; image size, 240 × 96 pixels as acquired during HS-AFM observations; frame rate, 3.3 frames/s.

**Movie S13.** HS-AFM movies of three representative examples of three RIG-I molecules bound to 5′ppp-dsRNA in the presence of 0.2 mM ATP and polyubiquitin on AP-mica. Magenta arrows indicate the 2CARD complex. Scan area, 100 × 80 $nm^2$; image size, 240 × 96 pixels as acquired during HS-AFM observations; frame rate, 3.3 frames/s.

**Movie S14.** HS-AFM movies of three representative examples of four RIG-I molecules (examples 1 and 2) and seven RIG-I molecules (example 3) bound to 5′ppp-dsRNA in the presence of 0.2 mM ATP and polyubiquitin on AP-mica. Magenta arrows indicate the 2CARD complex. Scan area, 100 × 80 $nm^2$ for examples 1 and 3, and 80 × 64 $nm^2$ for example 2; image size, 240 × 96 pixels as acquired during HS-AFM observations; frame rate, 3.3 frames/s.
